# Ethanol inhibits dorsomedial striatum acetylcholine release

**DOI:** 10.1101/2025.05.30.656893

**Authors:** Logan E. Slade, Charles C. Levy, Jacob Mitcham, Armando G. Salinas

## Abstract

Alcohol use disorder (AUD) has severe adverse health and economic impacts totaling over $240 billion annually. Despite this, FDA approved treatments for AUD are limited. For this reason, further understanding of the neurobiological mechanisms in AUD is required for new treatments. An aspect of AUD involves deficits in behavioral flexibility that is similarly seen in animal models with depletion of dorsal striatal cholinergic interneurons (CINs). We found that acute EtOH (40 mM) inhibits the firing rate of dorsal striatal CINs, which are the primary source of acetylcholine (ACh) in the dorsal striatum. Additionally, we found through slice photometry recordings using an intensity-based ACh sensing fluorescent reporter (iAChSnFR) that acute EtOH (40 mM) inhibits dorsal striatal ACh release. In accord, in vivo fiber photometry with iAChSnFR also showed inhibition of ACh release following acute EtOH (2 g/kg ip). To induce EtOH dependence in mice, we used the chronic intermittent EtOH (CIE) vapor exposure model. Following CIE, we found that CIE-treated mice had a significant depression of ACh release compared to control mice in the dorsomedial but not dorsolateral striatum. Then, we performed stereological cell counts of CINs in CIE and control mice to examine the cause of this ACh deficit and found that CIE mice had a significant decrease in CINs in the dorsomedial but not dorsolateral striatum. In conclusion, our data show that EtOH inhibits dorsal striatal cholinergic signaling in a subregion specific manner that may contribute to AUD related behaviors.

## Introduction

Alcohol use disorder (AUD) is a chronic brain condition affecting nearly 29 million Americans marked by a persistent inability to stop drinking despite adverse personal, professional, and health-related consequences [1,2]. This persistent alcohol use is linked to impaired cognitive flexibility and decision-making, traits believed to involve dysfunction in brain regions such as the dorsal striatum [3–7]. The dorsal striatum is involved in movement, motor control, learning, memory, habit formation, and appetitive behaviors, including those related to alcohol and substance use disorders [8–10]. The dorsal striatum can be functionally divided in rodents into dorsomedial (DMS) and dorsolateral striatum (DLS), with each subregion mediating different behaviors. For example, DMS is involved in goal directed behaviors while the DLS is involved in habitual behaviors [11–13]. As evidence grows supporting the role of DMS in mediating cognitive flexibility, understanding how chronic alcohol consumption disrupts this neural circuitry may be key to developing more effective treatments for AUD [14,15]. This is critical given the widespread impact of AUD, contributing to over 117,000 deaths annually and costing the U.S. >$240 billion each year [16,17].

Within the dorsal striatum, cholinergic interneurons (CINs) serve as the primary source of acetylcholine (ACh) and play a vital role in modulating striatal circuit activity, thereby influencing striatal output [18,19]. ACh signaling in the DMS has been specifically implicated in behavioral flexibility, and studies have highlighted CINs in the posterior DMS as critical contributors to this process, particularly in adapting behavior in response to salient or changing stimuli [20–22]. Mixed effects of EtOH on ACh release have been reported in various brain regions. For example, acute EtOH has been shown to decrease CIN firing rate and ACh release in striatal slices [23,24], but increase ACh release in the ventral tegmental area (VTA) and nucleus accumbens in vivo [25,26]. However, other studies suggest a dose dependent effect of EtOH on ACh release. Specifically, lower doses (<0.8 g/kg) of EtOH can increase extracellular ACh measured in vivo, while higher doses (>2.4 g/kg) were shown to decrease extracellular ACh [27]. Chronic EtOH has been shown to decrease ACh release measured in vivo in the prefrontal cortex (PFC) and hippocampus [28,29], but no results have been reported in the dorsal striatum. Chronic EtOH has also been shown to reduce cholinergic neurons in the basal forebrain of adult rats that were chronically treated during adolescence [30–32].

Given the mixed reports of EtOH effects on ACh signaling in the literature, our study seeks to address these discrepancies with real-time measurements of striatal ACh release both in vitro and in vivo. To do this, we used a novel genetically-encoded fluorescent biosensor, iAChSnFR, to measure real-time ACh release following acute and chronic EtOH treatment using a model of EtOH dependence that mirrors many of the facets of AUD. We employed electrophysiology, voltammetry, optogenetic, and immunofluorescent methods coupled with in vivo EtOH exposure and drinking assays to assess the effects of EtOH on striatal CINs and ACh release.

## Materials and Methods

### Subjects

Male C57BL6J mice (Jax, strain #000664, Bar Harbor, ME) were obtained at 9-10 weeks of age and acclimated at the LSUHSC Shreveport Animal Resources Facility for at least one week prior to any experiment onset. Body weights were collected at least once per week throughout each experiment. B6.129S-*Chat^tm1(cre)Lowl^*/MwarJ (ChAT-Cre) mice (Jax, strain #031661) and B6.Cg-*Gt(ROSA)26Sor^tm14(CAG-tdTomato)Hze^*/J (Ai14) mice (Jax, strain #007914) were obtained at 6-8 weeks of age and crossed to generate ChAT-Cre x Ai14 mice to allow for the visualization of CINs for the electrophysiological experiments.

Adult (>12 weeks) male and female mice were housed one to five per cage, with rodent chow and water available ad libitum, and were kept on a 12 h light/dark cycle. All procedures performed in this work follow the NIH guidelines and were approved by the LSUHSC-S and NIAAA Institutional Animal Care and Use Committees (protocols P-22-012 and LIN-DL-1, respectively).

### Chronic Intermittent Ethanol (CIE) Exposure

CIE exposure was induced via vapor inhalation as previously described [33–36] with minor modifications. Briefly, mice were exposed to EtOH vapor or room air for 16 hours per day starting at 17:00, followed by 8 hours of withdrawal starting at 9:00. This was repeated for four consecutive days (Monday-Friday), followed by a 72-hour withdrawal (Friday-Monday). This full week cycle was repeated four times. Mice did not receive an EtOH loading dose before the EtOH vapor exposure or pyrazole.

Custom designed alcohol vapor chambers were purchased from La Jolla Alcohol Research, Inc. (La Jolla, CA). EtOH is pumped into a heated flask and combined with an airstream to volatilize and carry the EtOH vapor into airtight chambers. The EtOH vapor concentration going into the chambers ranges from 2-3.5%. EtOH vapor levels are titrated to maintain BECs >200mg/dL on average. On occasion, some animals required subcutaneous fluids to aid in recovery.

### Ethanol Consumption

To confirm the CIE protocol induced EtOH dependence, a two-bottle choice (2BC) paradigm was used (Fig. 1A). Individually housed mice were given access to water and a 10% EtOH solution for 24 hours per day for 7 consecutive days (Monday-Monday) to establish baseline EtOH consumption. Mice were then randomly divided into two groups, CIE exposure and controls. The CIE protocol was conducted for two consecutive cycles, followed by a second week of 2BC. Then, 2 more CIE cycles were completed, followed by a third week of 2BC. Both bottles were weighed daily at the same time throughout the seven day 2BC periods. EtOH consumption was normalized to body weight (g/kg/day). Water consumption was also measured to determine water intake and EtOH preference. Additionally, total liquid intake was calculated by combing the consumption of both bottles in g/kg/day. Water and EtOH bottle positions were alternated daily to avoid a place preference.

**Figure 1.**
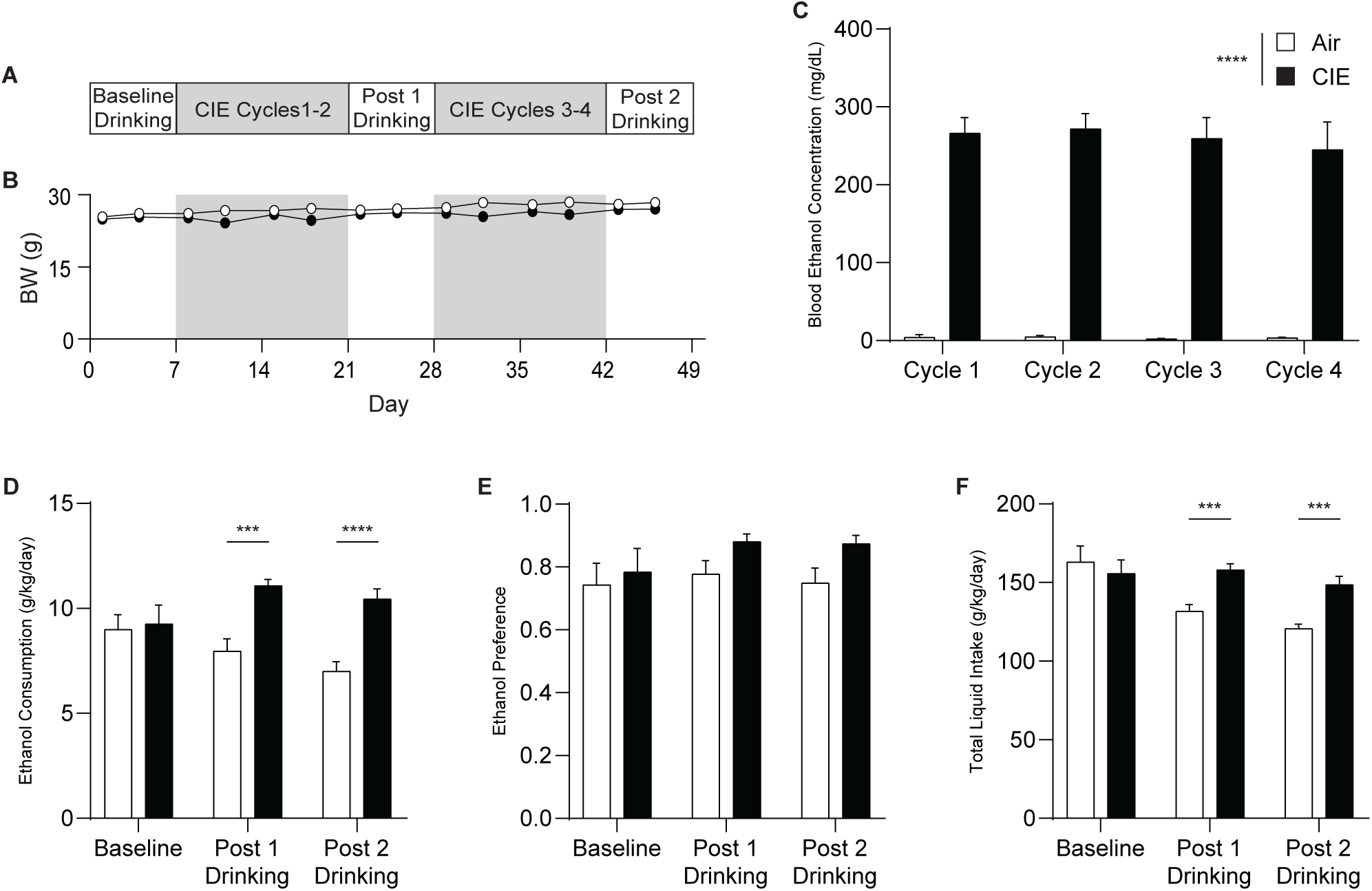
Chronic intermittent EtOH (CIE) exposure increases EtOH consumption. **A** Experimental timeline with **B** corresponding body weights averaged by treatment group. **C** BEC measurements taken during CIE. **D** EtOH consumption measured in g/kg/day via 2BC pre-CIE, post-CIE Cycle 2, and post-CIE Cycle 4. **E** EtOH preference measured to compare amount of EtOH consumed vs amount of water consumed during 2BC. **F** Total liquid intake combines EtOH and water consumption during 2BC. Air n=19 mice, CIE n=17 mice. ***p<0.001, ****p<0.0001. Error bars represent the SEM.

### Blood Ethanol Concentration (BEC) Measurement

BECs were determined each week from each animal undergoing CIE or control treatment using a commercially available assay (Pointe Scientific, Wayne County, MI) according to the manufacturer’s protocol with some modifications. Tail blood samples were collected the morning (9:00) after the second and fourth days of CIE exposure to ensure subjects were reaching BEC levels >200 mg/dL [34]. Blood samples were collected in heparinized capillary tubes (Fisher Scientific, Waltham, MA) and centrifuged at 3000rpm for 4 minutes in a hematocrit centrifuge (LW Scientific, MX12 Micro-Combo Centrifuge, Lawrenceville, GA). The plasma samples were then diluted 1:5 with water. Then 5 µLs of diluted plasma are added to 50 µL of alcohol reagent (Pointe Scientific, A7504) in a 96-well plate and incubated at 30°C for 5 minutes. After incubation, absorbance was measured in a microplate reader (Accuris Instruments, MR9600-T, Denver, CO) with a 340 nm filter. This absorbance reading was compared to a standard curve of known alcohol concentrations to determine the BEC for each subject. For brain slice photometry (BSP) and immunofluorescence experiments, BECs were determined once a week.

### Stereotaxic Injections

Subjects were bilaterally injected with the biosensor AAV9-EF1α-dbl flx-hChR2-H134R-eYFP (Addgene, 20298-AAV9, titer: 1x10^-13^ GC/mL, Watertown, MA) or AAV9-hSyn-iAChSnFR (Vigene Biosciences, titer: 1x10^-13^ GC/mL, Rockville, MD) for fast-scan cyclic voltammetry (FSCV) or photometry experiments, respectively, in the dorsal striatum (coordinates relative to bregma: +1.0 mm anterial, +/- 2.0 mm medial/lateral, -2.25 mm ventral). The virus was infused at a rate of 100 nL/min for a total of 500 nL per hemisphere. Subjects were allowed 2 weeks to recover before beginning CIE treatment.

For experiments comparing dorsal striatal subregions, injections were performed in the DMS (+1.0 mm anterior, +/-1.5 mm medial/lateral, -2.5 mm ventral) and DLS (+1.0 mm anterior, +/-2.25 mm medial/lateral, -2.5 mm ventral).

### Brain Slice Preparation

To prepare brain slices for electrophysiological and BSP experiments, subjects were anesthetized, rapidly decapitated, and brains extracted. Dorsal striatal slices were prepared at 225 µm or 300 µm thick (for electrophysiology and BSP/FSCV experiments, respectively) using a vibrating tissue slicer (Leica VT1200 S, Germany) in oxygenated cutting ACSF (containing in mM: 194 sucrose, 30 NaCl, 4.5 KCl, 26 NaHCO_3_, 1.2 NaH_2_PO_4_, 10 d-Glucose, 1 MgCl_2_, pH 7.4). Slices were then allowed to incubate in oxygenated recording ACSF (containing in mM: 124 NaCl, 4.5 KCl, 26 NaHCO_3_, 1.2 NaH_2_PO_4_, 10 d-Glucose, 1 MgCl_2_, 2 CaCl_2_, pH 7.4) for 30 minutes at 34 °C followed by a minimum of 60 minutes at room temperature. For voltammetry experiments, the recording ACSF recipe included, in mM: 126 NaCl, 2.5 KCl, 1.2 NaH_2_PO_4_, 1.2 MgCl_2_, 2.4 CaCl_2_, 0.4 L-ascorbate, 20 HEPES, 11 d-Glucose, 25 NaHCO_3_, pH 7.4. Expression of TdTomato (Ai14), ChR2-eYFP, or iAChSnFR was confirmed for the electrophysiology, FSCV, and BSP experiments, respectively, by imaging the acutely prepared brain slices with a Zeiss Axio Zoom.V16 microscope equipped with an X-Cite Xylis fluorescent lamp (Excelitas Technologies, XT720S, Pittsburgh, PA) Zeiss filter sets 43 and 38 (Ex: 545/25, Di 570, Em: 605/70 and Ex: 470/40, Di 495, Em: 525/50, respectively); Zeiss USA), and a Zeiss Axiocam 705 mono digital camera.

### Cell-attached Electrophysiology

Cell-attached electrophysiology was used to measure CIN firing rate in response to acute EtOH. ChAT-Cre x Ai14 mice were used and CINs were identified by the presence of TdTomato using a mCherry filter cube (Semrock BrightLine, LED-mCherry-A-OMF, Rochester, NY). A pipette filled with ACSF was used to create a tight seal (resistance >1GOhm) with the cell membrane. Cells were allowed to stabilize for at least 10 minutes before data collection. The baseline firing rate was determined from a 10 minute period in ACSF before switching to 40 mM EtOH for an additional 10 minutes, and followed by a 10 minute washout period in ACSF. **Fast-Scan Cyclic Voltammetry (FSCV)**

FSCV experiments were performed as previously described [37,38]. Briefly, striatal slices were moved to the rig and allowed to acclimate to the slice recording chamber for at least 10 minutes before beginning experiments. Voltammetry ACSF was continuously perfused on the slice at a rate of 1.5–2 mLs/min and at a temperature of 32 °C. Once ready a custom patch cord (FG400UMT, 400 µm, 0.39 NA, Thor Labs, Newton, NJ) with a flat cleave was lowered onto the slice into a region expressing ChR2-eYFP under a Zeiss Stereo Discovery.V12 microscope with an Excelitas X-Cite 120PC Q fluorescence lamp. The patch cord was coupled with a 470nm LED (M470F3, ThorLabs) to excite the ChR2-expressing CINs with a 2ms pulse width (4.6 µW/mm^2^). Then, a carbon fiber electrode (CFE) was placed into the slice in front of the fiber optic. Cyclic voltammograms were generated by applying a triangle waveform from −0.4 V to + 1.2 V to −0.4 V at a scan rate of 400 V/s every 100ms with a Chem-Clamp potentiostat (DAGAN) and DEMON Voltammetry and Analysis software [39]. CFEs were fabricated in house by aspirating a single, 7 µm diameter, carbon fiber strand (T650 fiber, Goodfellow, England) through a 1.2 mm OD x 0.68 mm ID borosilicate glass capillary tube (A-M Systems, Sequim, WA). The glass capillary tube was then placed in a P-1000 micropipette puller (Sutter Instruments, Novato, CA) and pulled to encase the carbon fiber in glass on one end. The pulled CFE was trimmed under 100X total magnification to a final exposed carbon fiber length between 150–250 µm past the glass seal.

For acute EtOH experiments, optical stimulation was applied every 5 minutes. Once responses became stable, four baseline measurements were collected before applying 50 mM EtOH onto the slice for 3 additional measurements. In a subset of slices, the nAChR antagonist mecamylamine was applied after EtOH treatment to confirm that the recorded dopamine (DA) responses were driven by ACh [40–42]. CFEs were calibrated against a 1 µM DA standard.

### Brain Slice Photometry (BSP)

BSP with iAChSnFR was used to assess evoked ACh release in response to electrical stimulation delivered with a constant current stimulus isolator (DigiTimer DS3, Fort Lauderdale, FL) and a bipolar stimulating electrode (Plastics One, Torrington, CT) placed within the fluorescing region on the surface of the slice. ACh release was evoked by a single electrical pulse (1.0 ms). In order to measure changes in florescence (ΔF/F), the recording region of interest (ROI) was determined using a camera (Thor Labs, CS165MU - Zelux 1.6 MP Monochrome CMOS) fixed to the microscope (Olympus, BX51WI, Center Valley, PA) at 4x magnification. Once a ROI was selected, magnification was increased to 40x to limit the sampling region to 160 µm x 120 µm region and measurements were collected with a PMT-based photometer. The microscope consisted of a photometer with an 814 photomultiplier (Horiba, PTI D-104, Irvine, CA) fixed with a GFP filter (Semrock BrightLine, GFP-4050A-OMF-ZERO) and a fluorescent lamp (Excelitas Technologies, X-Cite XYLIS LED XT720S). The microscope photometer output was digitized at 10 kHz with a Digidata1550B and data were analyzed offline with Clampfit11.2 software (Molecular Devices, San Jose, CA).

For acute EtOH treatment experiments, naïve C57 mice (>12 weeks old) were used. Dorsal striatal slices were prepared and BSP was performed to establish baseline changes in fluorescence at a constant stimulation intensity before switching to 40 mM EtOH for 15 minutes. Following EtOH application, we switched back to recording ACSF for an additional 15 minutes for the washout period. For CIE treated mice, subjects were sacrificed 3-5 days after the final CIE exposure and dorsal striatal slices were prepared. BSP was performed to collect Input-Output curves at various stimulation intensities (50-1600 µA every 3 minutes).

### In Vivo Fiber Photometry

In vivo fiber photometry was used to measure iAChSnFR activity in awake, freely-moving mice similar to our previous work [43]. After allowing at least 3 weeks for iAChSnFR or iAChSnFR NULL virus expression, a fiber optic probe (Thor Labs, CMFC22L05) was implanted. Following the fiber optic probe implant, mice were allowed to recover for at least one week before further experiments.

Photometric recordings were conducted using a custom-built photometry system previously developed in the lab [43]. This system included a 473 nm picosecond pulsed laser (BDL-473-SMC, Becker & Hickl, Germany) and a hybrid photo detector (HPM-100-40, Becker & Hickl). Samples were collected at 20Hz and hardware was controlled by a time correlated single photon counting module (SPC130-EM, Becker & Hickl) and SPCM software (v9.77, Becker & Hickl). A 3-meter, 200 µm core multimode fiber patch cord (FG200UCC; Thor Labs) terminating with a 2.5mm ceramic probe end was coupled to the implanted fiber optic probe on the mouse with a ceramic mating sleeve (ADAF1; Thor Labs). The signal from the iAChSnFR was passed through a 510/10 band pass emission filter (FF02-510/10, Semrock) before the hybrid detector. Mice were allowed to acclimate to the test room for one hour in their home cage while tethered to the photometry system. Then recordings began and a 20 minute baseline period was collected. Following this baseline period, mice were quickly injected with 2g/kg EtOH ip (20% w/v in saline), replaced into their homecages, and photometry recording resumed for one hour. Then mice were detached from the photometry system and returned to the mouse colony room. Data were analyzed offline and entered into Microsoft Excel for archiving and organizing. Data were then exported to GraphPad Prism 10 for graph preparation and Adobe Illustrator for figure preparation.

### Immunofluorescence (IF) Staining and Cell Counts

Following CIE or air exposure, subjects were anesthetized with isoflurane and transcardially perfused with a 4% paraformaldehyde/PBS solution (PFA). Brains were extracted and postfixed in PFA for 24 hours before being switched to PBS. The brains were then switched to a 30% sucrose/PBS solution for at least 48 hours before sectioning into 40µm slices in three series on a RM2125 RTS microtome (Leica) with a BFS-3MP freezing stage (Physitemp, Clifton, NJ). Sections were stored at -20C until IF labeling could be performed. Free floating sections were washed in PBS five times for 10 minutes, followed by two, one-minute washes in H_2_O, and two, 10-minute washes in 0.5% NaBH_4_/H_2_O. Then the sections were washed five times for 10 minutes in PBS, followed by a 30-minute wash in 0.2% Triton X-100/PBS (PBS-T) before a three-hour blocking step in 5% BSA/PBS-T. Sections were then incubated in a primary antibody solution consisting of Goat anti ChAT (1/500, Sigma AB144P, RRID: AB_2079751, St. Louis, MO) in 0.5%BSA/PBS-T at room temperature overnight. Sections were then washed four times, for 15 minutes in PBS. Sections were then incubated for 30 minutes in PBS-T, followed by incubation in a secondary antibody solution consisting of AlexaFluor568 donkey anti goat (1/2000, ThermoFisher Scientific A-11057, RRID: AB_2534104, Waltham, MA) at room temperature for three hours. After secondary antibody incubation, sections underwent five consecutive washes with PBS for 15 minutes before mounting and cover slipping with DAPI Fluoromount-G (SouthernBiotech, Birmingham, AL). Slides were sealed with clear nail polish and allowed to dry for at least two days before imaging.

Imaging of ChAT-ir neurons was completed using a Zeiss AxioZoom fluorescent microscope equipped with an X-Cite Xylis fluorescent lamp, Zeiss Filter Set 43, an Axiocam 705 mono digital camera, and Zen 3.5 software. Images were acquired at 23x magnification with 200x optical zoom for a total 4600x magnification at two focal planes per ROI. Two ROIs per striatal subregion were collected. For cell counts in ChAT-Cre x Ai14 samples, ChAT-ir+ cells were labeled using Rabbit anti DsRed (1/1,500) + Dk anti Rb 568 (1/1,000) and TdTomato+ cells were labeled using Goat anti ChAT (1/500) + Dk anti Gt 647 (1/1,000).

For stereological cell counts, the optical dissector method was used [44]. Images were loaded into ImageJ and the multipoint tool was used to perform cell counts. Counts were performed on all brain sections containing the dorsal striatum or pedunculopontine nucleus (PPN). Additional cholinergic nuclei were examined across other brain regions (Fig S1). For ChAT-Cre x Ai14 samples, stereological counting of cells was performed in each channel separately. The dorsal striatal sections were binned across the anterior-posterior axis into four or five points (for DMS and DLS, respectively). Then the mean cell number was determined. The total cell density was calculated as follows: 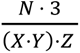, where *N* is the number of cells counted, *X* is the distance along the X-axis, *Y* is the distance along the Y-axis, and *Z* is the distance along the Z-axis of 3 slices (120 µm). Counts were performed by two observers and cross correlated to ensure accuracy. Counts from both observers were found to be within +/- 10% (Fig. S2).

### Statistical Analysis

For all BEC data, repeated measures Two Way ANOVAs were used with treatment group (Air or CIE) and CIE Cycle as the factors. For all 2BC data, repeated measures Two Way ANOVAs were used with treatment group (Air or CIE) as the between subjects factor and drinking stage (i.e. baseline, post-drinking 1, or post-drinking 2) as the within subjects factor with Sidak’s post hoc tests. For the effects of acute EtOH on CIN firing rate, FSCV, in vitro and in vivo ACh release, two-tailed paired t-tests were used. For ACh release following CIE, repeated measures Two Way ANOVAs were used with treatment group (Air or CIE) as the between subjects factor and stimulation intensity as the within subjects factor. For ChAT-ir+ cholinergic neuron counts of DMS and DLS, repeated measures Two Way ANOVAs were used with treatment group (Air or CIE) as the between subjects factor and distance from bregma as the within subjects factor. For ChAT-ir+ cholinergic neuron counts of PPN, a two-tailed t-test was used. For ChAT-Cre x Ai14 cholinergic neuron counts of DMS and DLS, repeated measures Two Way ANOVAs were used with treatment group (Air or CIE) as the between subjects factor and distance from bregma and channel as the within subjects factors. For ChAT-Cre x Ai14 cholinergic neuron counts of PPN, a two-tailed t-test was used.

## Results

### 2BC/CIE

Mice underwent a 2BC paradigm cycled with CIE exposure for a total of seven weeks. We found a main effect of CIE treatment such that CIE mice obtained higher BEC levels than air control mice controls (Fig. 1C, Two way ANOVA, F(1, 34)=303.9, p<0.0001). We found a main effect of CIE treatment such that CIE mice consumed more EtOH than air controls (Fig. 1D, Two way ANOVA, F(1, 34)=16.37, p=0.0003). Post hoc analysis revealed CIE mice showed significantly higher levels of EtOH drinking relative to air control mice at the Post-CIE 1 drinking stage (Sidak’s multiple comparisons test, t=4.65, p=0.0002) as well as the Post-CIE 2 drinking stage (Sidak’s multiple comparisons test, t=5.16, p<0.0001). We did not find a main effect of CIE treatment for EtOH preference (Fig. 1E, Two way ANOVA, F(1, 34)=3.62, p=0.066). We did, however, find a main effect of CIE treatment showing that CIE mice consumed more total liquid than air controls (Fig. 1F, Two way ANOVA, F(1, 34)=6.97, p=0.012). Post hoc analysis revealed CIE mice showed significantly higher levels of total liquid intake relative to air control mice at the Post-CIE 1 drinking stage (Sidak’s multiple comparisons test, t=4.7, p=0.0001) as well as the Post-CIE 2 drinking stage (Sidak’s multiple comparisons test, t=4.39, p=0.0004).

### Acute Ethanol Electrophysiology, FSCV, and iAChSnFR Photometry

To determine the effect of acute EtOH on CIN firing rate, we performed cell attached recordings and applied 40 mM EtOH. We found that acute EtOH decreased the average firing rate of CINs (Fig. 2D & E, Paired t-test, t=3.18, df=4, p=0.034). Further, while performing FSCV with optical stimulation, we found that acute EtOH (50 mM) decreased the ACh contribution to DA release (Fig. 2I & J, Paired t-test, t=5.22, df=15, p=0.0001). Application of the nAChR antagonist, mecamylamine (20 µM) completely blocked the optically-driven DA release confirming that the responses were due to release of ACh and mediated by nAChRs (Fig. 2H). In accord, we found that acute EtOH decreased evoked ACh release in the dorsal striatum (Fig. 2N & O, Paired t-test, t=2.96, df=4, p=0.042). Similarly, in vivo, we also found that acute EtOH depressed ACh release following a 2g/kg ip injection (Fig. 2S & T, Paired t-test, t=5.03, df=3, p=0.015).

**Figure 2.**
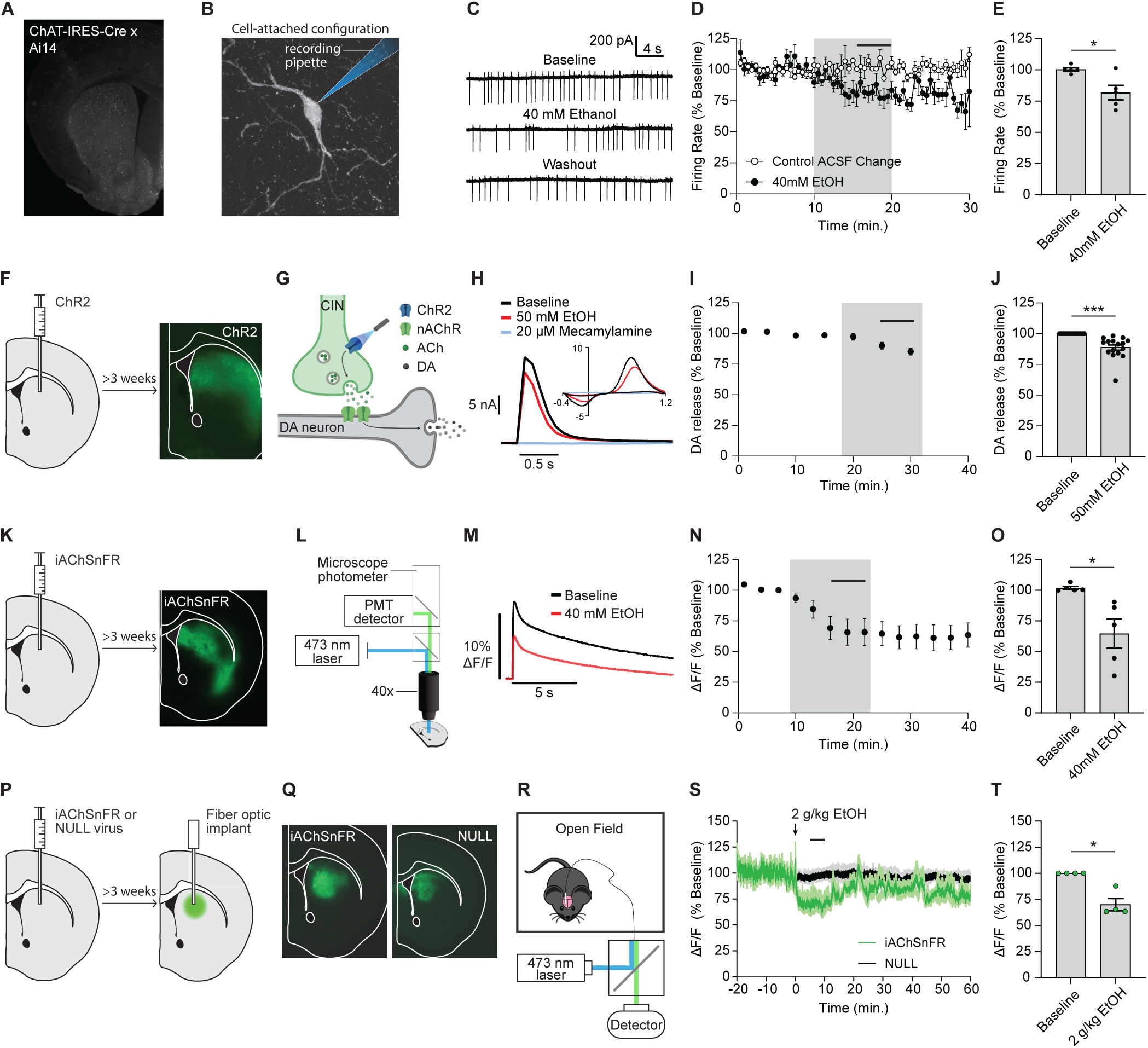
Acute EtOH effects on cholinergic signaling. **A** Fluorescent image of a striatal slice from a ChAT-Cre x Ai14 mouse. **B** CIN Cell-attached electrophysiology schematic. **C** Representative traces showing the acute effect of 40 mM EtOH on CIN firing rate. **D** Time course of 40mM EtOH (or ACSF control) effect on CIN firing rates (EtOH shaded region of graph). Control n=3 slices, EtOH n=5 slices. **E** Summary graph of 40 mM EtOH effect on CIN firing rate. **F** Schematic of ChR2 virus infusion in dorsal striatum and ChR2 expression. **G** Light activation of ChR2 causes ACh-driven DA release. **H** Representative traces showing the acute effect of 50 mM EtOH and 20 µM mecamylamine on ACh-driven DA release. **I** Time course of 50mM EtOH effect on ACh-driven DA release (EtOH shaded region of graph). n=16 slices. **J** Summary graph of 50mM EtOH effect on ACh-driven DA release. **K** Schematic of iAChSnFR virus infusion in dorsal striatum and iAChSnFR expression. **L** Microscope photometer setup for BSP recordings. **M** Representative traces showing the acute effect of 40 mM EtOH on evoked ACh release. **N** Time course of 40mM EtOH effect on evoked ACh release (EtOH shaded region of graph). n=5 slices. **O** Summary graph of 40mM EtOH effect on evoked ACh release. **P** Schematic of iAChSnFR (or NULL) virus infusions into dorsal striatum for in vivo fiber photometry experiments and fiber optic probe implant. **Q** Representative expression of iAChSnFR or NULL virus. **R** Schematic of open field fiber photometry setup for in vivo fiber photometry recordings. **S** Time course graph of ACh release following 2 g/kg EtOH (ip) administration. **T** Summary EtOH effect on in vivo dorsal striatal ACh release. *p<0.05, ***p<0.001. Error bars represent the SEM.

### Chronic Ethanol iAChSnFR Photometry

We found that CIE treated mice had significantly higher BECs than air control mice (Fig. 3C, Two way ANOVA, F(1, 16)=171, p<0.0001). Following CIE, we found that evoked ACh release was blunted relative to air controls (Fig. 3E, Two way ANOVA, F(1, 30)=5.49, p=0.026). We also found a main effect of stimulation intensity (F(1.69, 50.63)=240.5, p<0.0001). However, these recordings were done without regard to DMS-DLS regions. We, therefore, repeated these experiments with site-specific injections (i.e. DMS or DLS-targeted). In both the DMS and DLS, we found that CIE treated mice had significantly higher BECs than air control mice (Fig. 3G & K, Two way ANOVA, F(1, 8)=374.1, p<0.0001 and F(1, 8)=260.3, p<0.0001, respectively for DMS and DLS). We also found a significant main effect of stimulation intensity in both DMS (Two way ANOVA, F(1.84, 31.25)=211, p<0.0001) and DLS (Two way ANOVA, F(1.84, 31.3)=308.1, p<0.0001). In DMS we found that CIE treated mice had significantly less ACh release compared to air controls (Fig. 3I, Two way ANOVA, F(1, 17)=11.07, p=0.004). Post hoc analysis revealed CIE mice showed significantly less ACh release relative to air control mice at the 1600 µA stimulation intensity (Sidak’s multiple comparisons test, t=3.91, p=0.007). In the DLS, however, we found no significant differences in ACh release between treatment groups (Fig. 3M, Two way ANOVA, F(1, 17)=2.4, p=0.14).

**Figure 3.**
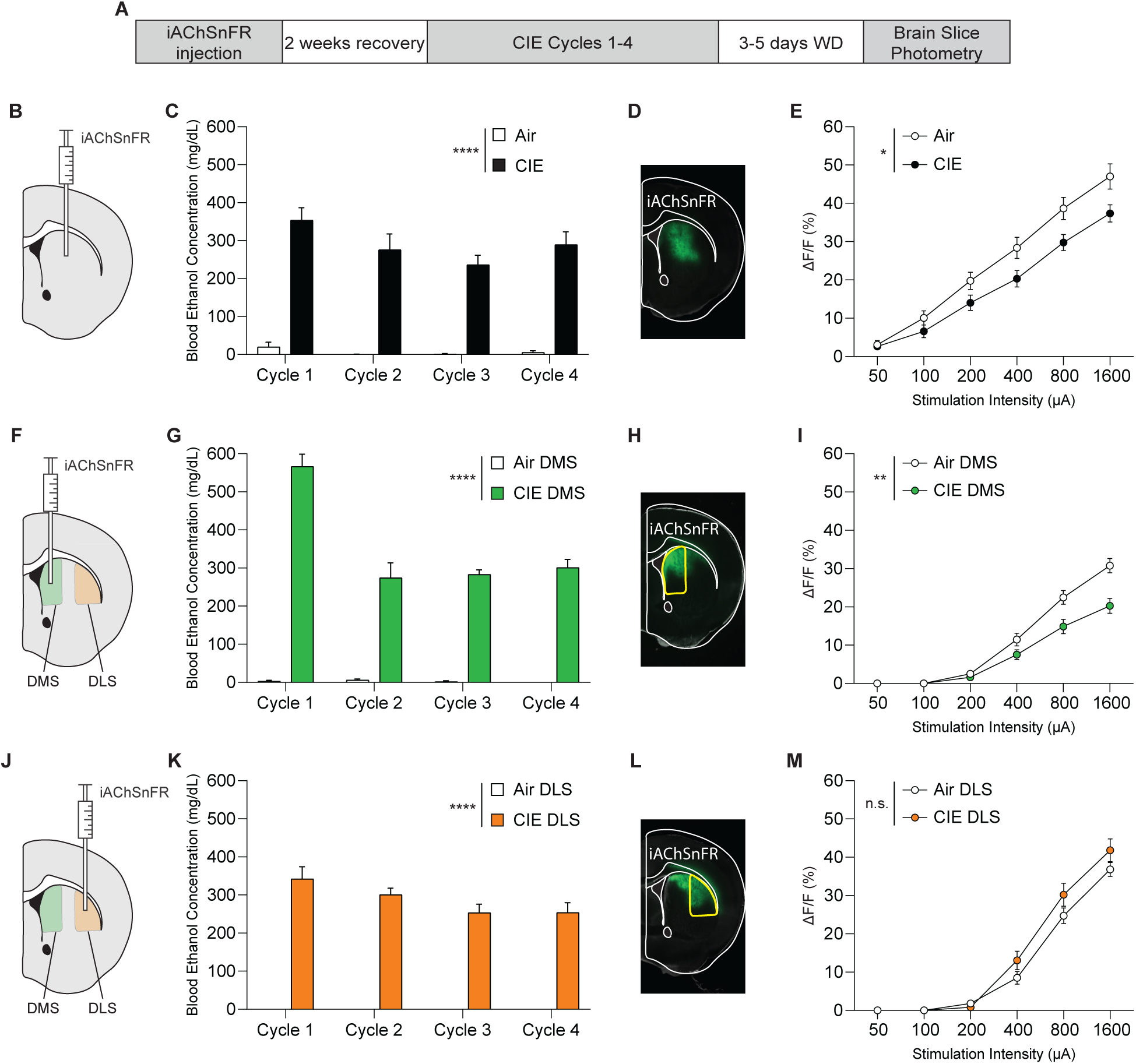
Chronic intermittent EtOH (CIE) exposure decreases evoked ACh release. **A** Experimental timeline. **B** Stereotaxic injection of iAChSnFR into dorsal striatum. **C** BEC measurements taken during CIE. Air n=10 mice, CIE n=8 mice. **D** Expression of iAChSnFR in dorsal striatum. **E** Input-Output curves examining the effect of stimulation intensity on ACh release in dorsal striatum. Air n=16 slices, CIE n=16 slices. **F** Stereotaxic injection of iAChSnFR into DMS. **G** BEC measurements taken during CIE. Air DMS n=5 mice, CIE DMS n=5 mice. **H** Expression of iAChSnFR in DMS. **I** Input-Output curves examining the effect of stimulation intensity on ACh release in DMS. Air DMS n=10 slices, CIE DMS n=9 slices. **J** Stereotaxic injection of iAChSnFR into DLS. **K** BEC measurements taken during CIE. Air DLS n=5 mice, CIE DLS n=5 mice. **L** Expression of iAChSnFR in DLS. **M** Input-Output curves examining the effect of stimulation intensity on ACh release in DLS. Air DLS n=11 slices, CIE DLS n=8 slices. *p<0.05, **p<0.01, ****p<0.0001. Error bars represent the SEM.

### Stereological Cell Counts

To assess whether the decreased evoked ACh release in the DMS following CIE was due to a loss of cholinergic neurons, we conducted ChAT IF labeling and stereological counting of cholinergic neurons in the dorsal striatum and PPN. For the DMS, a significant main effect of treatment was observed (Fig. 4E, Two way ANOVA, F(1, 19)=9.88, p=0.005), indicating a reduction in CINs following CIE exposure. Additionally, there was a main effect of distance from bregma (Fig. 4E, Two way ANOVA, F(3, 56)=45.87, p<0.0001). Post-hoc analyses using Sidak’s test revealed that the differences were most pronounced at +1.00 mm from bregma (p=0.02). There was no treatment × distance interaction (Fig. 4E, Two way ANOVA, F(3, 56)=0.36, p=0.78). In the DLS, no significant effect of treatment was found (Fig. 4F, Two way ANOVA, F(1, 19)=0.001, p=0.97), indicating that CIE did not affect CINs. There was a main effect of distance from bregma (Fig. 4F, Two way ANOVA, F(3, 55)=28.28, p<0.0001). There was no treatment × distance interaction (Fig. 4F, Two way ANOVA, F(4, 74)=1.95, p=0.11). In the PPN, no significant effect of treatment was found, indicating that CIE did not affect CINs (Fig. 4I, Unpaired t-test, t=0.6705, df=16, p=0.51).

**Figure 4.**
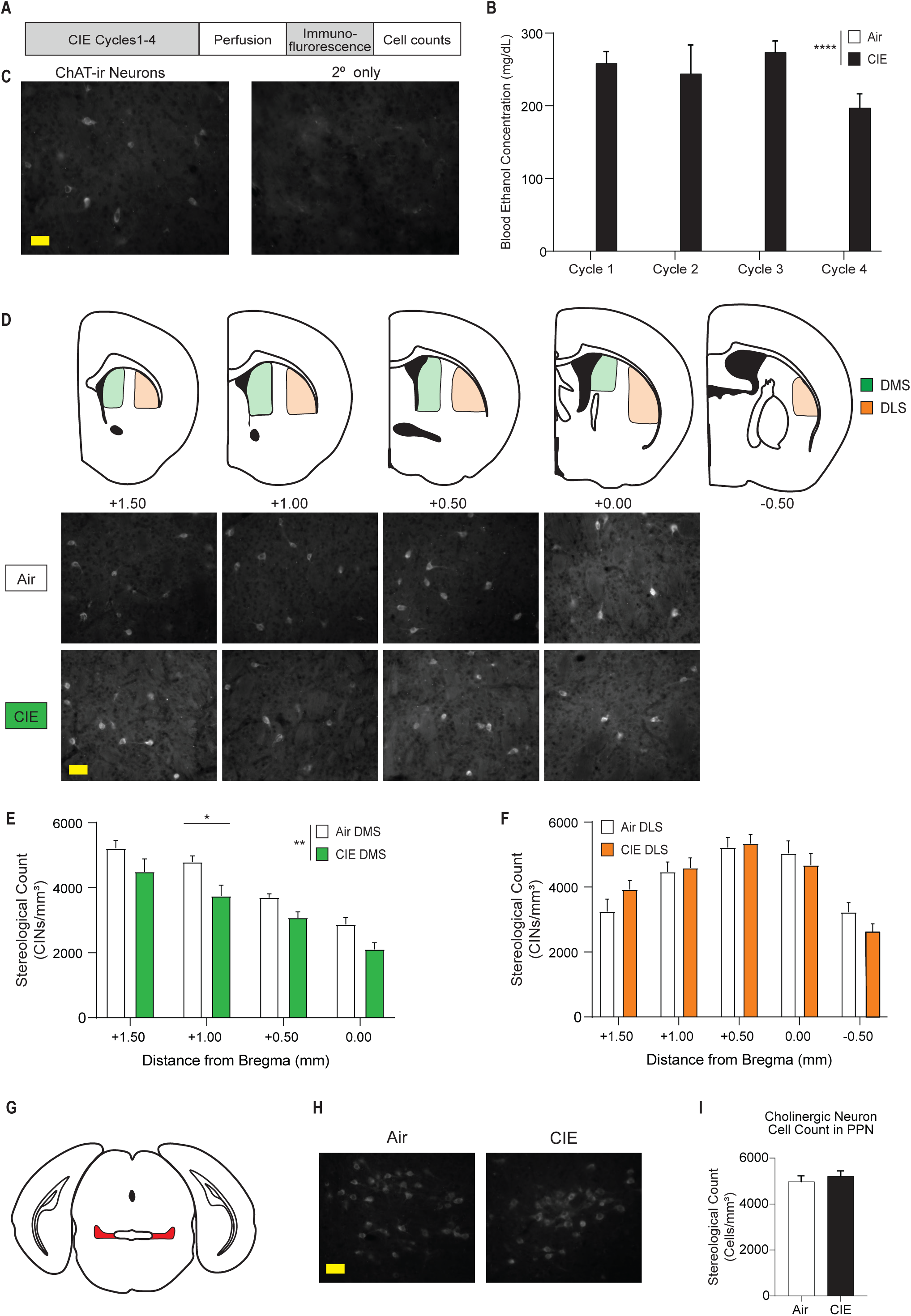
Chronic intermittent EtOH (CIE) treatment reduces DMS CINs. **A** Experimental timeline for stereological counting of CINs. **B** BECs were significantly elevated in CIE-exposed subjects compared to air controls. **C** Representative Choline Acetyltransferase (ChAT) IF labeling of CINs and secondary-only control. **D** Schematic showing the ROIs for stereological counting of CINs across various distances from bregma, with corresponding ChAT labeling images from the DMS. **E** CIE reduced ChAT-ir+ cells in the DMS but not **F** DLS. Air n=10 mice, CIE n=11 mice. **G** Schematic of ROI for stereological counting of PPN ChAT-ir+ cells. **H** Representative images of ChAT-ir+ cells in the PPN. **I** PPN ChAT-ir+ cells were similar between Air and CIE groups. Air n=8 mice, CIE n=10 mice. *p<0.05, **p<0.01, ****p<0.0001. Error bars represent the SEM. Scale bars=50μm.

To determine if the reduction in ChAT-ir+ cells was due to a decrease of ChAT expression or loss of CINs, we used ChAT-Cre x Ai14 mice in which ChAT neurons are permanently labeled regardless of changes to ChAT gene expression. For the DMS, a significant main effect of treatment was observed (Fig. 5D, Two way ANOVA, F(1, 5)=7.01, p=0.046), indicating a reduction in DMS CINs following CIE exposure. Additionally, there was a main effect of distance from bregma (Fig. 5D, Two way ANOVA, F(3, 13)=34.94, p<0.0001). Post-hoc analyses using Sidak’s test revealed that the differences were most pronounced at +1.00 and +0.50 mm from bregma (p=0.011, p=0.032, respectively). There was also a treatment × distance interaction (Fig. 5D, Two way ANOVA, F(3, 13)=4.94, p=0.018). In the DLS, no significant effect of treatment was found (Fig. 5E, Two way ANOVA, F(1, 5)=1.47, p=0.28), indicating that CIE did not affect DLS CIN counts. There was a main effect of distance from bregma (Fig. 5E, Two way ANOVA, F(2, 9.5)=31.38, p<0.0001). There was no treatment × distance interaction (Fig. 5E, Two way ANOVA, F(4, 19)=1.47, p=0.25). In the PPN, no significant effect of treatment was found, indicating that CIE did not affect CINs (Fig. 5F, Unpaired t-test, t=1.07, df=5, p=0.34).

**Figure 5.**
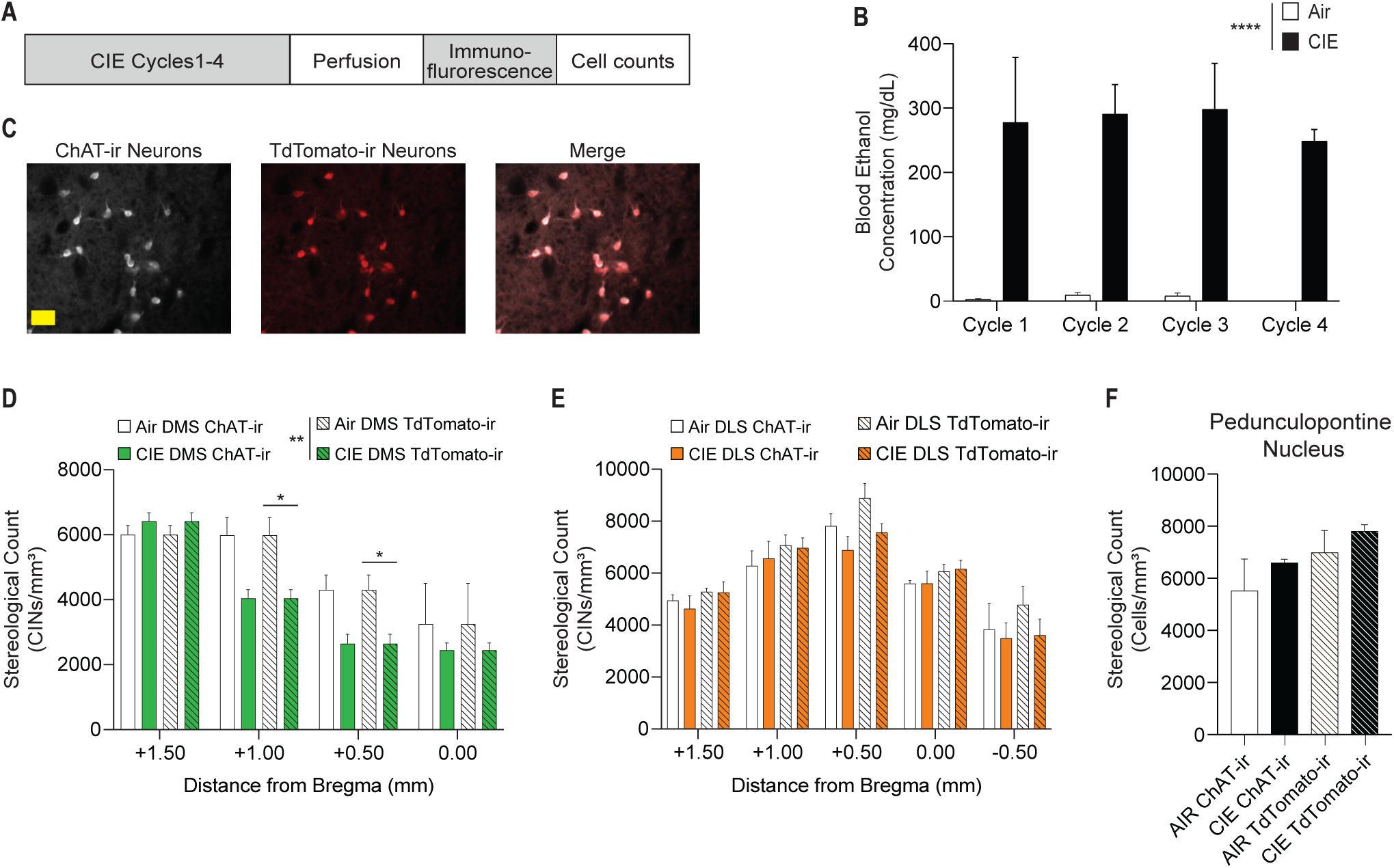
Chronic intermittent ethanol (CIE) treatment reduces DMS CINs. **A** Experimental timeline. **B** BECs were significantly elevated in CIE subjects compared to Air controls. **C** Representative Choline Acetyltransferase (ChAT) and TdTomato-ir IF labeling of CINs. **D** CIE reduced ChAT-Cre x Ai14 TdTomato-ir cells in the DMS but not **E** DLS. **F** PPN TdTomato-ir cells were similar between Air and CIE groups. Air n=3 mice, CIE n=4 mice. *p < 0.05. Error bars represent the SEM. Scale bar=50μm.

## Discussion

In this study, we found that acute and chronic EtOH inhibited dorsal striatal ACh release. Specifically, we found that chronic treatment resulted in depressed ACh release in the DMS but not DLS. Accordingly, we found a decrease in ChAT-ir+ neurons in the DMS but not DLS of chronic EtOH treated mice, which could underlie the decrease in ACh release observed.

The dorsal striatum has some of the highest levels of ACh in the brain [45–47]. The primary source of which is CINs, with a minor contribution from extra-striatal cholinergic neurons in the PPN [48–50]. Through cell-attached electrophysiological recordings, we observed a decrease in average firing rate of CINs following acute EtOH relative to basal firing rates. The mechanism driving this effect is undetermined but under investigation. Notably, this EtOH induced decrease in CIN firing rate matches previous findings from Blomeley, et al. ^23^. In that study, they found a 25% decrease of CIN firing rate following EtOH, which was similar to our findings. Notably, the Blomeley study performed electrophysiology using a perforated patch technique in rats while we used a cell-attached configuration in mice. Regardless, this EtOH induced decrease in CIN firing rate implies a subsequent decrease of striatal ACh release. To confirm this, we examined striatal ACh release, both directly and indirectly, with several methods (in vitro photometry, in vivo fiber photometry, and FSCV). We used optogenetics with FSCV to measure ACh-driven DA release and found that acute EtOH depressed ACh-driven DA release, suggesting that acute EtOH decreases ACh release. Similar to our FSCV findings, acute EtOH decreased dorsal striatal ACh release measured with iAChSnFR in vitro. This effect was also found in vivo using iAChSnFR with fiber photometry, further supporting our findings of EtOH induced inhibition of striatal ACh release through multiple techniques. This is the first real-time demonstration that EtOH inhibits dorsal striatal ACh release in vivo and in vitro. Notably, previous studies have shown EtOH induced increases in extracellular ACh in other brain regions like the VTA and nucleus accumbens using voluntary EtOH drinking paired with in vivo microdialysis or local reverse dialyzed EtOH microdialysis, respectively [25,26]. There are some notable differences between our study and the abovementioned studies including brain region, EtOH delivery method (slice application or ip injection vs drinking or perfused locally into the brain region of interest), and mode of ACh release measured (i.e. evoked release vs tonic release). Furthermore, other studies have revealed dose dependent effects of EtOH on ACh release with a lower dose increasing and a higher dose decreasing hippocampal ACh measured by in vivo microdialysis [27].

We next treated mice chronically with a variation of the CIE vapor exposure model of EtOH dependence that did not use an EtOH loading dose with pyrazole. This treatment still led to enhanced EtOH consumption in CIE subjects relative to controls. Our first in vitro photometry experiments found a significant decrease in ACh release in CIE treated subjects relative to controls. Earlier work performed by Kipp and Savage ^29^ showed similar results. In their study, rats underwent adolescent intermittent ethanol (AIE) exposure and ACh release in the PFC was measured in vivo using GRAB ACh 3.0 with fiber photometry. Their results showed a deficit in ACh release in response to cue and reward in the AIE treated rats relative to controls.

We followed up our in vitro experiments performing IF stereotaxic cell counts of CINs. Unexpectedly, we found that CINs were decreased in a striatal subregion dependent manner which prompted us to re-examine our BSP data with regard to striatal subregions. A cursory analysis suggested a striatal subregion specific deficit in ACh release in CIE treated subjects (data not shown). We followed up on this by performing DMS or DLS specific iAChSnFR viral infusions. In these mice, we found that CIE treatment decreased ACh release in the DMS but not DLS, mirroring our IF cell counts. We next considered whether the differences in IF cell counts could be due to chronic EtOH induced decreases in ChAT expression or loss of CINs. To address this, we used ChAT-Cre x Ai14 mice in which the TdTomato in these mice is driven by the constitutively active CAG promoter. Thus, following recombination by Cre, ChAT neurons would be permanently labeled, regardless of changes to ChAT gene (or protein) expression. Similar to our IF cell counts in C57BL6/J mice, ChAT-Cre x Ai14 mice treated with CIE had decreased DMS but not DLS CINs, suggesting that chronic EtOH is specifically pernicious to DMS CINs and that this loss of CINs may underlie or contribute to the decreased ACh release we observed in our BSP studies.

Studies utilizing AIE have shown that AIE treated rats had fewer ChAT-ir+ cholinergic neurons in the adolescent basal forebrain and into adulthood [30–32]. In addition to the loss of ChAT-ir+ cholinergic neurons, AIE was shown to reduce other cholinergic markers such as the high-affinity nerve growth factor receptor tropomyosin receptor kinase A (TrkA) and the low-affinity NGF receptor p75^NTR^ [30]. In accord, chronic EtOH consumption was shown to decrease ChAT activity as well as high-affinity choline uptake in the frontal cortex, nucleus basalis of Meynert, and parietal cortex [51]. These findings, in conjunction with ours, offer support for a neurotoxic effect of EtOH on cholinergic neurons. Interestingly from our study, the EtOH-induced decrease of ACh release and loss of CINs was only seen in DMS, suggesting a fundamental difference in toxicity pathways between striatal subregions and, possibly, the existence of multiple CIN subtypes. This idea is supported by previous reports identifying CIN subtypes that differed according to their expression of LHX6 [52]. Supporting this notion, previous studies have found that serotonin modulated CINs in a striatal subregion specific manner [53,54]. Specifically, the work performed by Virk, et al. ^54^ showed that bath application of serotonin (30 µM) decreases CIN firing rate in the ventral striatum but increases CIN firing rate in the dorsal striatum. This finding represents possible mechanistic differences of CINs located in the ventral and dorsal striatum which could explain the ACh release difference between previous reports in the nucleus accumbens [26] and our data in the dorsal striatum.

Our observed dorsal striatal subregion difference may contribute to the enhanced EtOH consumption we observed. The relative decrease in cholinergic transmission in the DMS, which is critical for the acquisition of goal-directed learning, may bias animals towards more habitual behaviors like EtOH consumption. DMS CINs have been demonstrated to play a role in the reversal of instrumental learning [12,21]. Thus, the loss of these CINs following chronic EtOH may hinder an animal’s ability to stop consuming EtOH. Supporting this, a recent study showed that chronic EtOH drinking in mice reduces thalamic excitatory inputs onto CINs in the DMS and impairs the reversal of action-outcome contingency, indicating reduced behavioral inflexibility [14]. This decrease in thalamic drive onto CINs could also contribute to the selective decrease in ACh release in the DMS we found following CIE treatment.

Overall, our findings demonstrate that both acute and chronic EtOH exposure inhibit ACh signaling in a subregion-specific manner within the dorsal striatum. This inhibition was due to the selective loss of CINs in the DMS, indicating that DMS CINs are particularly susceptible to the effects of chronic EtOH. These results suggest the existence of a distinct, region-specific CIN subpopulation within dorsal striatum. Future research will focus on comparing the vulnerability to chronic EtOH of CINs in the DMS and DLS, which could pave the way for targeted therapies aimed at minimizing or preventing EtOH toxicity in dorsal striatal cholinergic signaling and associated increases in EtOH seeking behaviors.

## Data Availability Statement

The data that support the findings of this study are available at lsuhs.figshare.com or by request from the corresponding author.

## Acknowledgements

Figure 2G: Created in BioRender. Slade, L. (2025) https://BioRender.com/kl0k7n2

## Author Contributions

This study was conceived and designed by AGS. AGS and LES drafted and revised the manuscript. LES performed experiments. LES collected and analyzed data. CCL and JM collected and analyzed cell count data. LES and CCL created figures. All authors reviewed the article and approved the submitted version.

